# A somatic evolutionary model of the dynamics of aneuploid cells during hematopoietic reconstitution

**DOI:** 10.1101/355826

**Authors:** Andrii I. Rozhok, Rebecca E. Silberman, Angelika Amon, James DeGregori

## Abstract

Aneuploidy is associated with many cancers. Recent studies demonstrate that in the hematopoietic stem and progenitor cell (HSPC) compartment aneuploid cells have reduced fitness and are efficiently purged from the bone marrow. However, early phases of hematopoietic reconstitution following bone marrow transplantation provide awindow of opportunity whereby aneuploid cells rise in frequency, only to decline to basal levels thereafter. Here we demonstrate by Monte Carlo modeling that two mechanisms could underlie this aneuploidy peak: rapid expansion of the engrafted HSPC population and bone marrow microenvironment degradation caused by pre-transplantation radiation treatment. Both mechanisms reduce the strength of purifying selection acting in early post-transplantation bone marrow. We explore the contribution of other factors such as alterations in cell division rates that affect the strength of purifying selection, the balance of drift and selection imposed by the HSPC population size, and the mutationselectionbalance dependent on the rate of aneuploidy generation per cell division. We propose a somatic evolutionary model for the dynamics of cells with aneuploidy or other fitness-reducing mutations during hematopoietic reconstitution following bone marrowtransplantation.

**Significance:** Bone marrow transplantations (BMT) following ablative irradiation pose a great health risk. It’s been shown that additionally the bone microenvironment is conducive to elevated frequencies of aneuploid cells in mice during bone marrow reconstitution post-BMT. As aneuploidy is linked with many cancers, we explore the reasons of such aberrant cell frequency peaks by Monte Carlo modeling. We demonstrate that elevated rates of aneuploidy early post-BMT are likely to be caused by reduced purifying somatic selection resulting from the expansion of the reconstituting population and the damage to stem cell niches caused by ablative radiation

## Introduction

Aneuploidy, or deviation of the chromosome number from the normal karyotype (resulting from chromosome mis-segregation), is associated with many cancers, being prevalent in both solid cancers and leukemia (1–4). The effects of aneuploidy vary, with some cellular phenotypes dependent on what specific chromosome(s) is lost or gained, and other phenotypes arising from a general stress response to aneuploidy (5). Consequently, associations of aneuploidy with cancers range widely, from a few percent, such as the loss of chromosome 1 or gain of chromosome 5 in kidney adenocarcinoma, to 50%, such as the loss of chromosome 3 in melanoma, and even 70% for the loss of chromosome 22 in meningiomas (2). In total, almost 90% of cancers exhibit gains or losses of at least one chromosome arm, with patterns specific to particular tumor types (4). For example, squamous cell cancers originating in multiple organs exhibit a common pattern of chromosome arm 3p loss and 3q gain.

Aneuploid cells have been shown to drive adaptation in yeast (6–8). This evidence has led to speculations that by conferring greater adaptability, variability of chromosomal ploidy in a cell population might lead to the expansion of aneuploid clones in human tissues, fueling further accumulation of oncogenic alterations in cells and progression to cancer (9–11). However, aneuploidy has been shown to more commonly reduce the fitness of animal somatic cells (5), just as most chromosome gains in yeast reduce their fitness (12, 13). Similarly, mice engineered to model the human trisomy of chromosome 21, the cause of Down syndrome, demonstrate lower proliferative potential of hematopoietic stem cells (HSC), mammary epithelial cells, neural progenitors and fibroblasts (14). Multiple mouse models of spindle assembly checkpoint mutants are lethal indicating that high level chromosome mis-segregation is highly detrimental (15). Aneuploidy has also been shown to promote premature differentiation and depletion of neural and intestinal stem cells in *Drosophila melanogaster* (16). We recently examined how aneuploidy impacts hematopoietic stem and progenitor cell (HSPC) fitness using transplantation of bone marrow from aneuploid or aneuploid-prone mouse models (15). This study demonstrated that increased aneuploidy is associated with reduced somatic stem cell fitness *in vivo*, as such aneuploid cells are efficiently purged from the hematopoietic compartment.

These experiments raise the question of how aneuploidy can be so tightly associated with a vast array of cancers, given that cancer development requires a series of expansions and fitness gains by more proliferative cell clones. One answer would be that only specific types of aneuploidy are involved in cancer. However, evidence shows that aneuploidy has various degrees of association with cancers across the board, including a gain or loss of almost any human chromosome (4). We performed computational modeling that indicates that rapid expansion of the engrafted HSC population together with reduced support of HSC stemness from damaged bone marrow microenvironments are plausibly the two primary mechanisms weakening purifying selection in early post-transplant bone marrow, providing a window of opportunity for the expansion of aneuploid HSCs. These results have implications for the generation of aneuploid cells in other contexts, including during cancer development.

## Results

In the context of bone marrow transplantation in mice, we previously showed that the peripheral blood descendants of aneuploidy-prone HSPCs demonstrate an immediate and substantial rise in the frequency of aneuploidy after bone marrow transplantation, despite a clear fitness disadvantage relative to euploid cells (15). For these experiments, aneuploid cells were generated at an increased rate due to a hypomorphic mutation in the mitotic spindle assembly checkpoint protein gene BUB1-related 1 (*BUBR1*). As shown in **Fig. 1A**, following the rise in the fraction of aneuploid cells in peripheral blood post-transplantation, the frequency of aneuploid cells subsequently declines to the low baseline levels typical of unperturbed blood cells (15). Given that peripheral blood is regularly generated from HSC and downstream progenitors in the bone marrow, this pattern suggests that early reconstitution provides a window of opportunity for the enrichment of aneuploid self-renewing cell types, such as HSC and HSPC, despite their lower fitness. We currently do not have data to discriminate whether the observed aneuploidy originates from HSC or later more committed cell lineages of hematopoiesis. However, as aneuploidy declines later during BM reconstitution, we focus this research on finding the answer to the observation that tolerance of aneuploidy demonstrates a temporal change, being higher early in the process. Thus, while our model starts with parameters determined for HSC, the general principles to be explored here should be relevant for different progenitor stages and cell lineages.

**Fig. 1.**
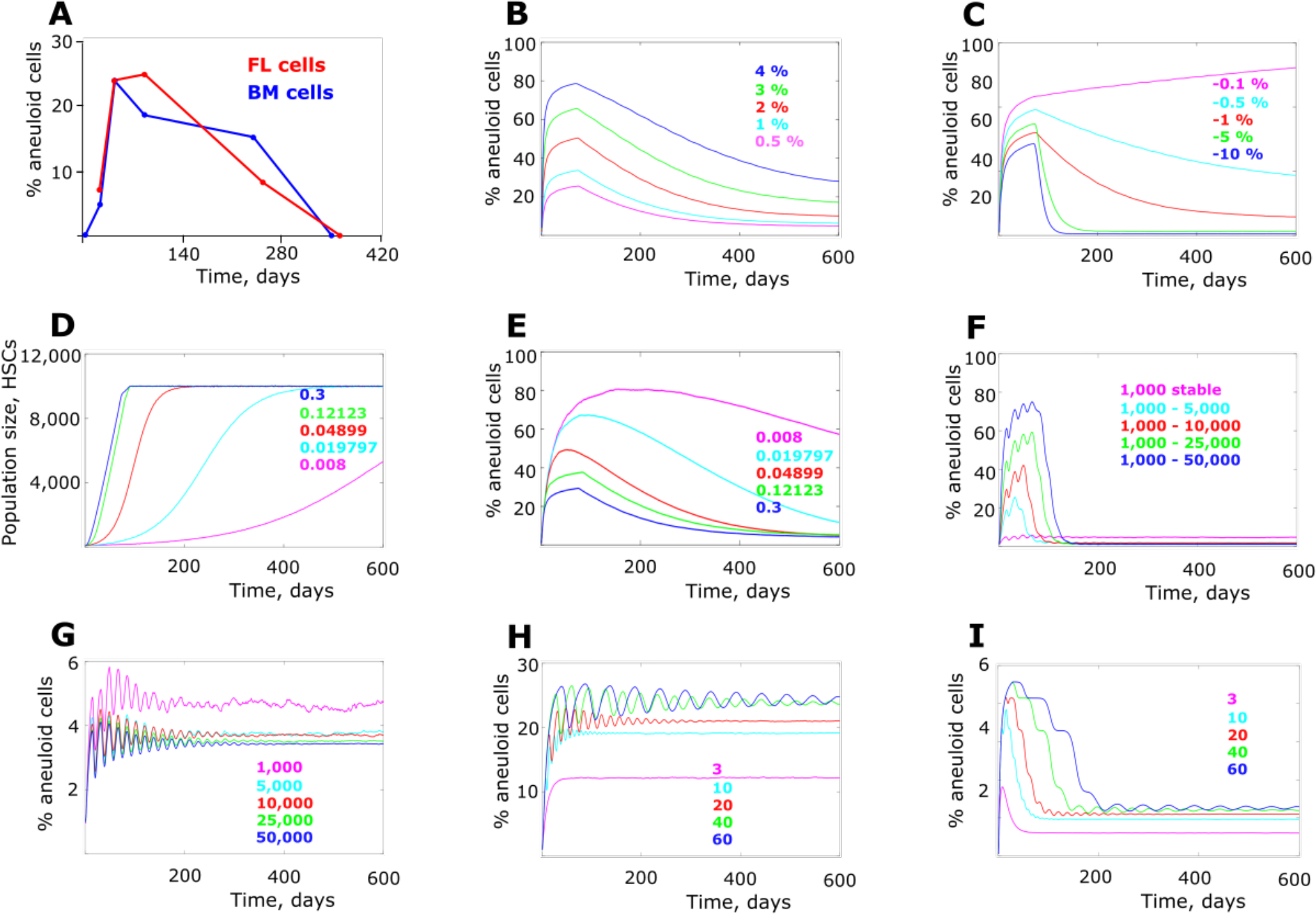
Frequency of aneuploid HSC cells in the simulated post-transplantation bone marrow. **(A)** Frequency of aneuploid cells observed in peripheral blood of recipient mice after receiving transplanted bone marrow from BUBR1^H/H^ (hypomorphic) mice; data are from Pfau et al. (15); data at days 350 and 364 says was collected following the protocol in Pfau et al. (15); FL-transplanted fetal liver cells, BM - transplanted bone marrow cells; see Supplements section Aneuploidy counts for a summary of data. **(B)** Simulated aneuploidy dynamics with varying aneuploidy generation rate per cell division (numbers color-matched to respective data lines; statistics in **Fig.S2**). **(C)** Simulated aneuploidy dynamics with a range of cell fitness cost induced by aneuploidy (statistics in **Fig.S3**). **(D)** Dynamics of HSC population increase post transplantation over time (color-matched numbers represent growth coefficients which determined the shape of the population size growth). **(E)** Simulated aneuploidy dynamics under various cell population expansion regimens (numbers color-matched as in (D); statistics in **Fig.S4**). **(F)** Simulated aneuploidy frequency at stable cell division rate of 1 in 20 days and various extent of cell population size expansion (color-matched numbers indicate initial and final population size in # of cells; statistics in **Fig.S5**). **(G)** Simulated aneuploidy frequency at a stable cell division rate of 1 in 20 days and different stable cell population sizes (color-matched numbers indicate population size in # of cells; statistics in **Fig.S6**). **(H)** Simulated aneuploidy frequency at a stable population size of 10,000 cells and varying stable cell division rates (color-matched numbers indicate the average interval in days between successive cell divisions; statistics in **Fig.S7**). **(I)** Simulated aneuploidy frequency under population expansion from 1,000 to 10,000 cells and varying stable cell division rates (color-matched numbers as in (H); statistics in **Fig.S8**).

The early phase of bone marrow reconstitution after transplantation differs from steady-state hematopoiesis in several respects. First, HSCs and hematopoietic progenitors are known to divide much faster immediately after transplantation and return to their normal cell cycle rate later (17). Early post-transplantation bone marrow also has free niche space after irradiation kills recipient HSCs, such that the transplanted population is not at an equilibrium but expands until the entire bone marrow niche space is reclaimed by the engrafted HSCs, in order to restore normal hematopoiesis. We expect that other progenitor compartments will behave similarly, as their numbers are reduced post-irradiation followed by recovery. Also, radiation exposure causes substantial damage to the bone marrow microenvironment, including via genomic damage, oxidative stress causing profound inflammation (18, 19). Such perturbation likely reduces the functionality of HSC niches for stem cell maintenance, similarly to the effect shown for mesenchymal stem cells (18). Niche perturbation should thus reduce the strength of purifying selection by impairing the support of HSPC relatively independently of HSPC phenotype.

### Model architecture

Using Matlab, we simulated bone marrow reconstitution by creating a virtual niche space as a matrix of 10,000 single-cell niches, based on estimated numbers of HSC in mice (20, 21). The initial number of HSCs was 100, reflecting the approximate number of HSC in a million transplanted mouse bone marrow cells (15). We set a rate of generation to aneuploidy and the average fitness cost relative to normal HSC and explored a range of both parameters in simulations. Cell division rate started from fast division (once per ^~^3 days) and returned to once per ^~^40 days when bone marrow HSC population size returned to the physiological ^~^10,000 cells. Total reconstitution was achieved in ^~^8 simulated weeks. The simulation lasted for 600 simulated days with daily updates. At each update, HSCs divided stochastically based on the current division rate. Excess cells were removed as a result of a binomial trial with probabilities of staying an HSC based on the number of HSCs after division, the assumed current population size based on the growth curve, and their relative fitness.

HSC niches had an additional property - the ability to maintain HSC stemness dependent on niche health after irradiation. This ability was realized by implementing an additional binomial “survival” trial with a certain probability for an HSC to leave the pool regardless of the cell’s relative fitness. Percentages of aneuploid cells were then tracked over the entire simulation time (600 days) under various values of model parameters. Whenever not directly manipulated as shown in **Fig. 1B,C** and **Fig. 2D,E**, aneuploidy generation rate per cell division was kept at 1%, and aneuploid cell fitness effects at −1% as standard parameters for all experiments. Higher generation rates and fitness effects are also explored. Cell division rate profile was as described in **Methods** and shown in **Fig. S1** whenever such a parameter was not varied on purpose or unless indicated otherwise. More details of the model are explained in **Methods** and the Matlab code is provided in **Supplements, section “Model code”**. The niche health model and rationale will be explained later.

**Fig. 2.**
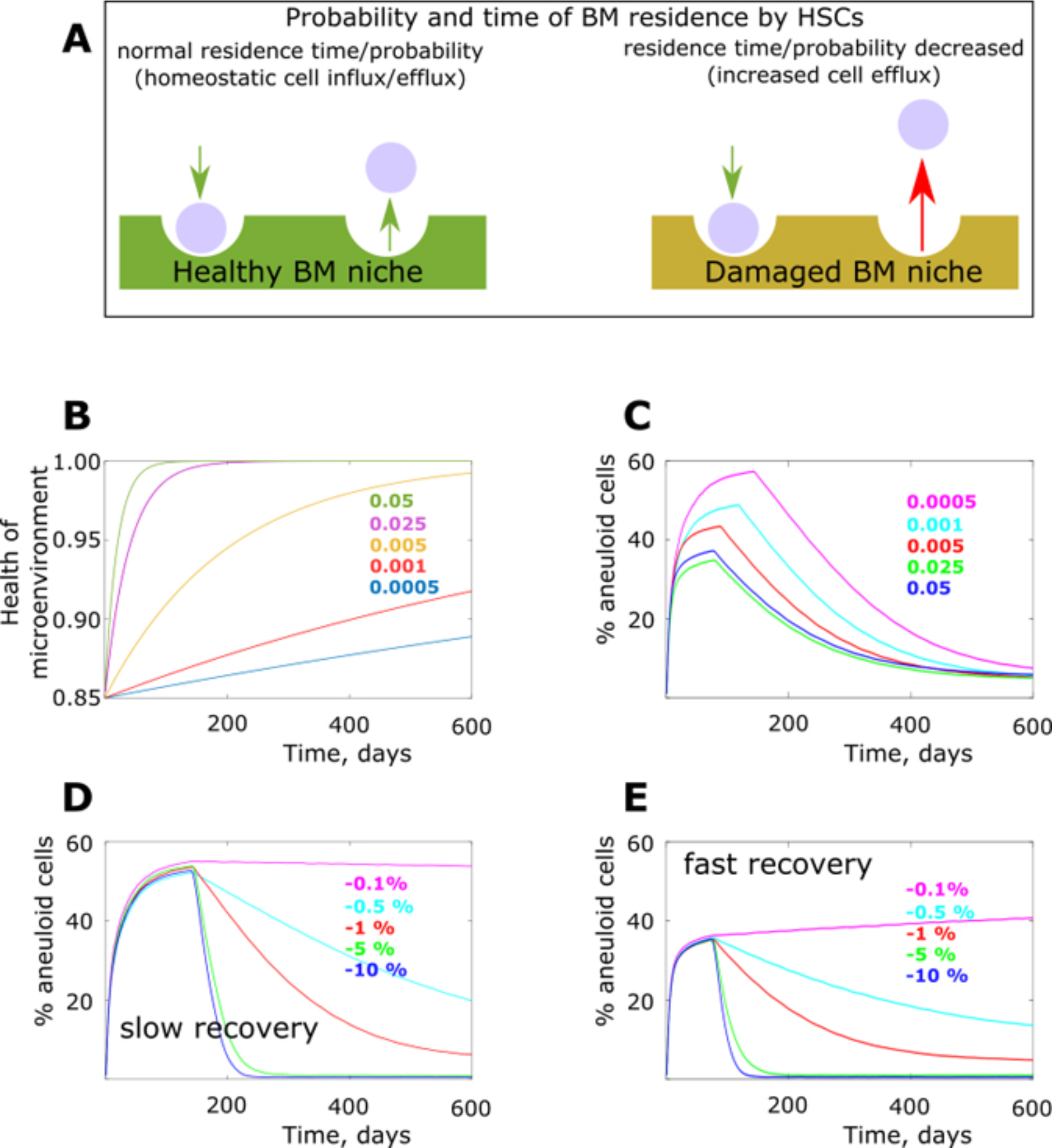
The model of degraded bone marrow niches and their effect on the aneuploidy dynamics. **(A)** Damaged bone marrow niche model (BM stands for bone marrow). **(B)** Temporal profiles of post-radiation bone marrow niche healing; Y at X=0 represents initial bone marrow niche health as a fraction of the maximum health equal to 1; color-matched numbers represent coefficients of healing speed in the function of bone marrow niche health of time; the curve with the coefficient 0.005 was used as standard in simulations were this parameter was not investigated. **(C)** Simulated aneuploidy dynamics under various profiles of post-radiation bone marrow healing; numbers (healing temporal profile coefficients) indicate as in (B) but color-matched separately (statistics in **Fig.S9**). **(D)** Simulated aneuploidy dynamics with a range of aneuploidy cell fitness cost (color matched numbers) and a slow bone marrow healing profile (coeff=0.0005, according to (B); statistics in **Fig.S10**). **(E)** Simulated aneuploidy as in (D) but under a rapid bone marrow healing profile (coeff=0.05, according to (B); statistics in **Fig.S11**).

### Niche-independent effects of bone marrow transplantation on aneuploid cell frequency

We set out to explore if altered purifying selection underlies the observed aneuploidy peak following bone marrow transplantation (see **Fig. 1A**). As shown in **Fig. 1B**, with a fitness cost applied to aneuploid cells relative to normal cells, the model replicated the general pattern observed experimentally, with aneuploid cell frequency peaking early and subsiding afterwards. Varying aneuploidy generation rate per cell division with a fixed aneuploidy fitness cost of −1% significantly affected the height of the frequency peak, while the pattern of the subsequent frequency decline remained similar (**Fig. 1B**). This result indicates that the mutation-selection balance that depends on the aneuploidy generation frequency per cell division is one of the evolutionary forces that should impact the aneuploidy frequency peak, particularly its height. We further set a fixed aneuploidy generation rate at 1% and applied the following range of aneuploidy fitness costs [−10%, −5%, −1%, −0.5%, −0.1%]. While increasing fitness cost resulted in a lowering of the peak of aneuploidy (**Fig. 1C**), the effect was relatively modest over the 100-fold range of fitness costs, indicating that purifying selection is weak during the expansion phase. In contrast, fitness cost demonstrates a significantly stronger effect on the purging of aneuploid cells that follows this peak, consistent with a strengthening of purifying selection during the post-reconstitution period. **Fig. 1C** also shows that if the aneuploidy fitness cost is low enough, aneuploidy frequency does not decrease. These results demonstrate that one explanation for the post-transplantation frequency of aneuploidy that we have observed earlier (15) could be the shifting balance between mutation (aneuploidy generation) rate and the strength of purifying selection.

Pre-transplantation radiation treatment eliminates resident HSCs and thus vacates HSC niche space for transplants, leaving room for the transplanted HSC population to expand. Population expansion should increase the presence of random drift and reduce the strength of purifying selection (22–24). As shown in **Fig. 1D**, we applied a range of HSC population expansion rates following transplantation with the same maximum population size. The rate of population expansion demonstrates an inverse relationship with the peak frequency of aneuploidy (**Fig. 1E**), with faster expansion lowering the aneuploidy peak. A number of confounding factors could potentially interfere with the effects of population expansion. First, a more rapidly increasing population size can counteract the effect of population expansion, whereby the expansion itself relaxes selection while the resulting increased population size intensifies it. Also, the rapidly dividing HSCs during the early post-transplantation phase could also intensify the strength of selection by increasing the number of cell generations per time unit, as argued previously (25).

To isolate the effect of population expansion per se and explore the effect of these additional factors, we first fixed the simulated cell division rate to an intermediate average of 1 division per 20 days throughout the simulation run. We further explored a range of population expansion rates from 1,000 cells to [1,000; 5,000; 10,000, 25,000; 50,000]. As shown in **Fig. 1F**, without the contribution of the changing cell division rate, greater population expansion caused higher aneuploidy frequency peaks. The effect of the selection-intensifying cell population size was still present, but obviously overcome by the selection-reducing effect of population expansion. Interestingly, with no expansion the aneuploidy peak was absent, validating a role of population expansion in generating the peak (**Fig. 1F**). In order to test the effect of population size we tested the model with a range of stable population sizes [1,000; 5,000; 10,000; 25,000; 50,000] and with the same stable cell division rate as in **Fig. 1F**. In the absence of the effects of population expansion and the changing cell division rate, we see that population size does have the predicted selection-suppressing effect, producing differences in peak aneuploidy frequency early and influencing the frequency during later phases of reconstitution. However, the difference was tangible only between the smallest population size (1000 HSC) and the rest, revealing non-linear effects and indicating that at the size of 5,000 and above drift is perhaps a minor factor (**Fig. 1G**). However, in the absence of the early-phase population expansion, HSCs do not demonstrate an early frequency peak.

We further fixed population size at 10,000 cells and applied a range of stable cell division rates at once per [3, 10, 20, 40, 60] days to isolate the effect of cell division rates. As predicted, **Fig. 1H** Demonstrates that faster cell division does have a suppressive effect on aneuploidy frequency, presumably by intensifying the strength of selection (by increasing the number of cell generations per unit time). Just as in **Fig. 1G**, an aneuploidy peak is not produced in the absence of population expansion. In order to corroborate the role of expansion per se, we further applied population expansion from 1,000 to 10,000 cells but across the same range of fixed cell division rates at once per [3, 10, 20, 40, 60] days. **Fig. 1I** Demonstrates that in the presence of population expansion the model generates an early peak of aneuploidy frequency. Interestingly, the height of the peak and the rate of the subsequent aneuploidy elimination is dependent on cell division rate, with faster rates being more aneuploidy-suppressive, consistent with the results in **Fig. 1H**. Based on these results, we can conclude that HSC population expansion during early post-transplantation bone marrow reconstitution is likely to have a profound effect on the intensity of purifying selection and is another likely mechanism underlying the pattern observed *in vivo* (15).

Combined the results shown thus far indicate that the early post-transplantation peak of aneuploidy could be explained by shifts in the character of purifying selection that occur during hematopoietic reconstitution.

### Effects of bone marrow niche/microenvironment on aneuploid cell frequency

Another process characteristic of bone marrow transplantation following ablation of the resident bone marrow cells with radiation is damage to and recovery of the bone marrow microenvironment. The post-radiation bone marrow microenvironment appears to be highly perturbed by direct radiation-induced damage and the ensuing reactive oxygen species generation and inflammation (18, 19). Similar to such effects shown for mesenchymal stem cells (18), these changes should reduce the ability of bone marrow niches to support stemness in HSC, which require proper microenvironmental signaling to maintain homeostatic differentiation rates (26). We therefore further reasoned that the perturbed microenvironment should decrease the ability of HSC to maintain stemness, and this influence should be less discriminating between aneuploid and normal HSCs, being poorly supportive for all HSC (**Fig. 2A**). Such an effect might reduce the strength of selection based on intrinsic cell fitness differences.

In order to model the damaged niche shown in **Fig. 2A**, we added a stochastic effect exerted by the bone marrow niche and affecting all cells by adding an additional small probability within each cell’s binomial trial that the cell will leave the pool, as shown in the increased “cell efflux” model in **Fig. 2A**. This effect was then reduced over time following a certain function and reflecting the bone marrow healing process. A range of “healing” functions was used, with a healing coefficient 0.005 (**Fig. 2B**) used as standard in simulations where bone marrow recovery function was fixed. The function’s initial value (Y-axis in **Fig. 2B** at X=1) reflects the initial probability of a cell to maintain stemness per trial (lower probability indicates a more degraded niche). In other words, simulations started with the presence of an elevated HSC efflux from the niche (independent of cell phenotype) caused by bone marrow damage, and this effect gradually subsided over time as bone marrow “healed”.

In simulations with the niche effect added, we found that the default aneuploidy fitness cost of −10% was more appropriate to use, because the added aneuploidy-promoting effect from niche degradation drove aneuploidy peaks to higher frequencies compared to niche-independent modeling, often up to the point of fixation. Notably, the real fitness cost of aneuploidy in HSC is not known and is likely distributed depending on various types of aneuploidy. This fact, however, does not confound the investigation of the general principles underlying the observed aneuploidy peak. **Fig. 2C** Shows that the dynamics of niche healing has a significant effect on the aneuploid cell frequency peak and it also affects the time when the peak is reached. Slow bone marrow healing promotes an increased frequency of aneuploidy cells, showing that damaged bone marrow could counteract purifying selection relative to healthy bone marrow. To further investigate the ability of bone marrow health to impact purifying selection acting on aneuploid cells, we applied a range of aneuploidy fitness costs as indicated previously [−10%, −5%, −1%, −0.5%, −0.1%], and tested the system under two regimens of niche recovery: slow (coefficient of healing 0.0005 in **Fig. 2B**) and fast (coefficient of healing 0.05). Comparison of the resulting dynamics demonstrates that the early increase and the maximal aneuploidy frequency reached are essentially unaffected by the aneuploidy fitness cost in the presence of the damaged bone marrow microenvironment, consistent with very weak purifying selection. Notably, in the absence of such bone marrow damage effects, altering the fitness cost of aneuploidy does affect the height of the aneuploidy peak, albeit not proportionally to change in fitness (**Fig. 1C**).

In contrast, during the later phase of aneuploidy frequency reduction, where purifying selection is supposed to intensify when the HSC population has stabilized and the bone marrow microenvironment is healthier, we observe a substantial effect of fitness (**Fig. 2D,E**). After peaking, the frequency of aneuploid cells decreases dependent on the fitness cost of aneuploidy, and the effect of varying microenvironment health is minimal. Comparison of **Fig. 2D** and **Fig. 2E** further shows that the rate of bone marrow microenvironmental recovery has a significant effect on the height <u>and</u> timing of the peak frequency of aneuploidy, whereby slow niche recovery promotes aneuploidy and delays the onset of the second (aneuploidy reduction) phase.

We further explored the interaction of population expansion (according to the scheme in **Fig. 1D**) and bone marrow microenvironmental health by simulating the initial bone marrow health of 90% (relatively good) and 70% (more degraded), both under the healing coefficient of 0.005 (see **Fig. 2B**). Comparison of **Fig. 3A** and **Fig. 3B** Demonstrates that in a relatively healthy bone marrow microenvironment (**Fig. 3A**), the population expansion effect is present, but peak aneuploidy cell frequency is significantly lower than in a degraded bone marrow microenvironment (**Fig. 3B**). This result demonstrates a profound effect of bone marrow microenvironment degradation in suppressing purifying selection and promoting aneuploidy. More profound damage (**Fig. 3B**) to bone marrow also reduces the population expansion effect.

**Fig. 3.**
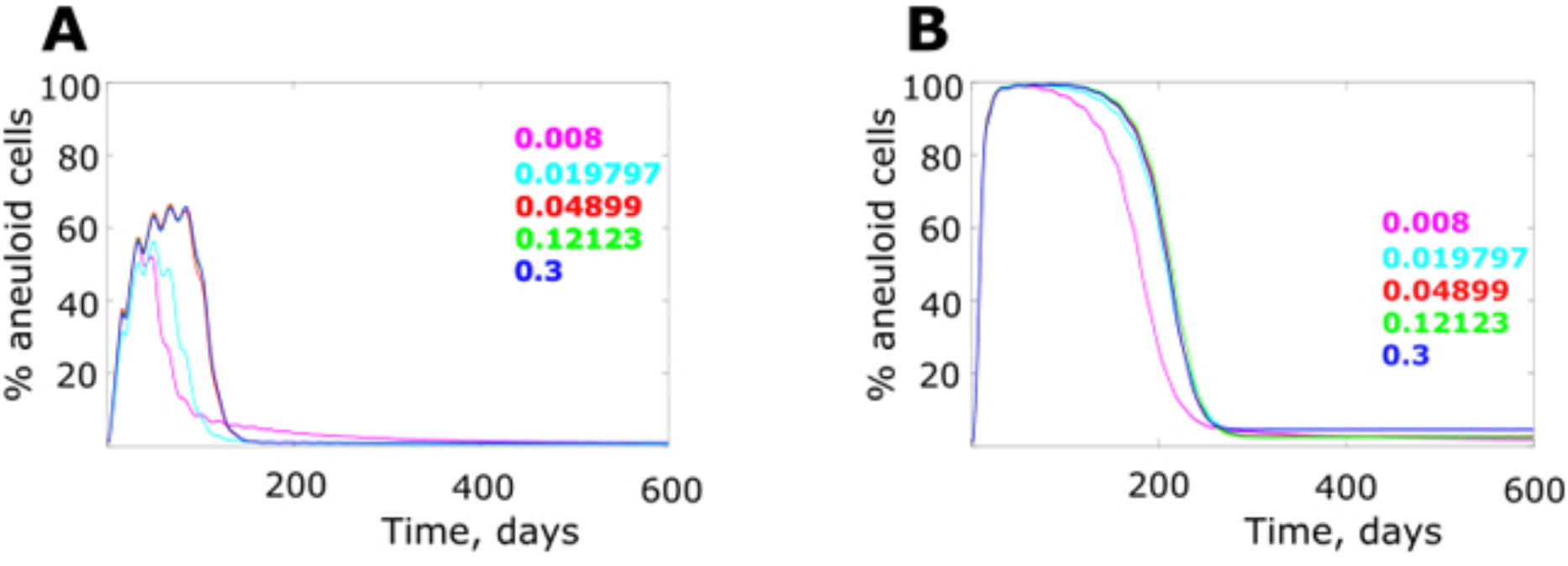
The effect of bone marrow health and population expansion on simulated aneuploidy dynamics. **(A)** Relatively healthy bone marrow (initial niche health 90%; healing profile coeff=0.005, according to Fig. 2B; statistics in **Fig.S12**). **(B)** Relatively degraded bone marrow (initial niche health 70%; healing profile as in (A); statistics in **Fig.S13**).

## Discussion

Our results demonstrate that the pattern of aneuploidy frequency during post-transplantation hematopoietic reconstitution observed by Pfau et al. (15) is likely the result of relaxed purifying selection during the early post-transplantation period. We demonstrate that at least two mechanisms could promote the observed aneuploidy frequency peak. First, a rapid expansion of the engrafted HSC population results in a reduction in the strength of somatic purifying selection by introducing an increased presence of drift, resembling the pattern shown in general population biology studies (22–24). Additionally, pre-transplantation ablative radiation treatment, by damaging and perturbing the bone marrow microenvironment, should degrade the capability of the niche to maintain HSC stemness, which we propose should affect all HSC with reduced or minimal discrimination of cell phenotypes. Our results demonstrate that such niche damage should promote an increase in the frequency of aneuploid cells and that this effect can be strong enough to overcome the contribution of population expansion or the fitness cost of aneuploidy. Recovery of the bone marrow microenvironment over time restores the power of purifying selection, leading to elimination of aneuploid cells from the pool. Our results also demonstrate, consistent with general population biology, that increased cell population numbers act to suppress aneuploidy by reducing the amount of drift and elevating thus the strength of purifying selection. Rapid cell division rates early post transplantation have the same effect, intensifying selection by increasing the number of cell generations per time unit.

We propose a model to explain changes in aneuploid HSC frequency post-transplantation (**Fig. 4**). The proposed scenario is that early hematopoietic reconstitution provides a window of opportunity for mutant cell clones of lower fitness, such as the aneuploid cells modeled here. This window is created by exposing transplanted HSCs to conditions of reduced strength of purifying selection. Later, purifying selection regains strength as the bone marrow microenvironment heals and the engrafted HSC population stabilizes at its homeostatic size, leading to elimination of mutant cells from the pool.

**Fig. 4.**
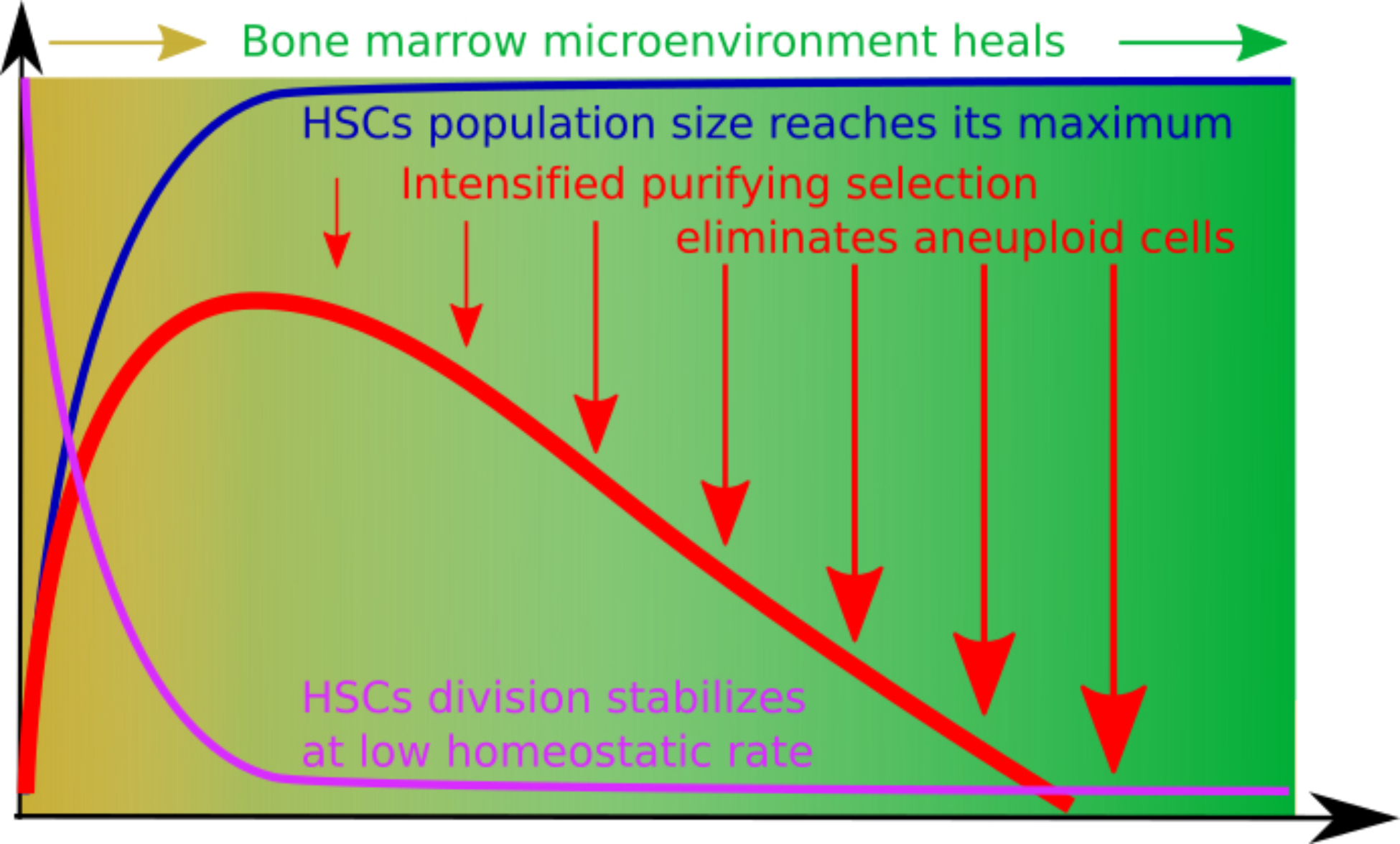
A model of factors influencing the strength of purifying selection in the post-radiation post-transplantation bone marrow. Early post-transplantation bone marrow is characterized by a highly perturbed microenvironment, as well as rapid HSC population expansion. This combination of factors reduces the strength of purifying selection and promotes drift. Later, HSC population numbers reach their maximum and the bone marrow microenvironment partially restores from radiation damage, processes that intensify the strength of purifying selection and lead to the elimination of aneuploid HSCs from the pool. The rapid cell division rates and the increased HSC population size (per se, excluding the effect of expansion) should act to suppress aneuploidy frequency.

The observations made by Pfau et al. (15) and the mechanisms we propose to explain them may have important implications for cancer research. First of all, bone marrow transplantation is known to be associated with a higher post-transplantation risk of leukemia (27). This increased risk has been attributed to the compromised immune system (28) and radiation-induced mutations (29–31). Another mechanism that could contribute, in line with our results, would be conditions of reduced purifying selection that are conducive to greater proliferation of pre-malignant mutant cells contained in the engraft bone marrow due to the perturbed bone marrow microenvironment and cell population expansion.

We have argued earlier that a similar decreased purifying selection might be involved in childhood leukemia (32). Although fetal development likely proceeds in a homeostatic environment, HSC population expands during development, which should promote a greater role for random drift, providing a window of opportunity for the accumulation and expansion of pre-malignant mutant clones that later in life could be purged by purifying selection. Interestingly, the increased aneuploidy observed during fetal development, which is subsequently purged (33), could potentially be similarly explained by relaxing purifying selection associated with rapid tissue growth (although typically with healthy tissue niches). That damaged microenvironments could weaken purifying selection could also be relevant for tissue in the elderly. Increases in expanded hematopoietic clones are relatively common in the elderly (34–36). While we and others have proposed that these expansions result from increased positive selection for mutations that confer adaptation to the aged bone marrow microenvironment (37, 38), results presented here further indicate that relaxed purifying selection in the degraded bone marrow microenvironment could contribute to such somatic evolution by being more permissive to phenotypic variability. However, aging-related changes likely do not provide a level of acute damage to bone marrow comparable to radiation treatment, but instead likely promote gradual alterations of the bone marrow microenvironment.

In each case, a temporary tolerance of aneuploidy (due to refilling of niches or due to degradation of niches) should increase the chances for a cell with a chromosomal aneuploidy event that would normally reduce fitness to undergo additional chromosomal rearrangements or other mutations that could decrease the cost of the initial aneuploidy. The more expanded the original aneuploidy cells are, the greater the odds of subsequent compensating events, the selective pressure for which should increase as the strength of selection returns post-bone marrow recovery. These chromosomal reassortments could generate more diversity in the population for potential oncogenic selection.

In a growing tumor, the disruption of normal niches could similarly contribute to the increased presence of aneuploidy, by reducing the cost. From this perspective, the presence of aneuploidy in a cancer can be interpreted as not simply an increased rate of chromosomal gains and losses (“genomic instability”), but as changes in the strength of selection that alter the frequency of aneuploidy cells *independent* of their generation rate. Relaxed purifying selection in cancers, as previously shown (39–42), could even create permissive conditions for cells with large-scale genomic perturbations, such as chromothripsis (43).

The modeling results presented here should stimulate further experimental efforts to decipher the somatic evolutionary processes and mechanisms that govern stem cell dynamics under conditions of rapid population expansion and/or damaged microenvironments.

## Methods

Simulations were performed in the Matlab programming environment (MathWorks Inc., MA). The model incorporates a matrix of simulated cells that divide with an average specified frequency determined by the curve of cell division shown in **Fig. S1**. Cells started from the average cell division rate of once per 3 days and reached the finals rate of once per 40 days (20, 21), with the dynamics of change corresponding to the dynamics of population size (the closer to the final size the slower the cells became). Cell division is stochastic and determined by a normal distribution with the mean frequency determined by the curve in **Fig. S1** and standard deviation equal to mean/8, as used in (44). The cell population matrix had a time-specific maximum capacity (cell number) which is determined by the population growth curves as shown in **Fig. 1D**. The simulation continued for 600 time units (days) and was updated at each time unit.

At each cell division, aneuploid cells were generated with the probability of aneuploidy per cell division as shown in **Fig. 1B** (1% in most simulations, unless indicated otherwise). Euploid cells were assigned fitness equal to 1. Aneuploid cells were assigned fitness as shown in **Fig. 1C** (−1%, or 0.99, in most simulations unless indicated otherwise). Based on cell fitness, the current matrix capacity and the total number of cells after cell divisions, at each simulation update each cell had a probability of leaving the pool (simulating death or differentiation) weighed by their fitness so that the remaining number of HSCs approximately corresponded to the current matrix capacity (fitness based competition for limited space). This probability was realized in corresponding binomial trials for each cell at each simulation update. The effect of niche degradation was realized by adding additional probability (in a binomial trial) for each cell to leave the pool; this probability did not depend on a cell’s fitness, replicating a hypothesized effect of phenotype-indiscriminate lower capacity of bone marrow niche to support stemness. The probability was proportional to niche health (the Y-axis value at X=1 in a chart exemplified by **Fig. 2B**). For example, if the initial bone marrow niche was considered 15% degraded, each cell had an additional probability of 0.15 of leaving the pool at each simulation update. This effect was reduced over time, following the healing curves shown in **Fig. 2B**, with a corresponding decrease in this probability as the function approached 1 over time (perfect niche, all competition is based exclusively on intrinsic HSC fitness), reflecting bone marrow niche health recovery with time past transplantation.

Simulations started with the initial population size of 100 HSCs, according to data from (15). All resulting aneuploidy curves are averages of 100 simulation repeats for each condition. The Matlab code for the model is presented in Supplements, section “Model code”.

Statistical comparisons of the simulated clonal dynamics were performed using the Matlab Statistics toolbox. Each simulated condition was run in 100 repeats. In order to elucidate as much statistical information about the relative behavior of clones as possible, we applied the following statistical procedure. At each time point (out of the 600 total simulation time points), we compared different conditions each represented by a sample of 100 runs by the Kruskal-Wallis method, which is a non-parametric version of ANOVA. The obtained p-values were plotted along the X-axis (simulation time points), with the Y-axis representing p-values (see Fig. S2-S13). This procedure allows visualizing the temporal dynamics of the differences in clonal behavior. The general magnitude of the difference in clonal behavior over time in this way can be visualized by the total sum of p-values (area under the p-value curve). We calculated this area and divided it by the total area of the chart, the latter being 1×600. The total area represents a hypothetical scenario whereby p-values are equal to 1 during an entire simulation, meaning that the compared behavior of clones was identical throughout the simulation. Respectively, if the area under the p-value curve equals zero, it would mean that such clonal behaviors are totally distinct throughout the simulation time. Realistically, however, p-values always are within that range and never reach such extremes. Therefore, the above-mentioned ratio shown in the top right corner of the chart in Fig. S2-S13, reflects the overall relative magnitude of the difference in clonal behavior throughout the compared simulations. The smaller the ratio, the greater the overall difference in clonal behavior. Following this statistical procedure, thus, we can demonstrate both the significance of the difference at each time point (p-value curve) and the overall magnitude of the difference throughout the simulation time.

Hematopoietic reconstitutions were performed following the protocol in Pfau *et al.* (2016). Briefly, B6.SJL-Ptprc^a^Pepc^b^/BoyJ (CD45.1) female mice were purchased from Jackson Laboratory and served as recipients for all reconstitutions. At 6-8 weeks old, recipients were treated with 9.5 Gy in a single dose, administered via 137^Cs^ irradiator (γ cell 40) at a dose rate of ^~^ 100 cGy/min, and were intravenously injected with 10^6^ donor (CD45.2) cells. In bone marrow reconstitutions, donor bone marrow cells were isolated from a BubR1^H/H^ donor at 5-7 weeks old. Red blood cells were lysed in ACK lysing buffer. The remaining white blood cells were counted on a Cellometer Auto T4 automated hemacytometer (Nexelcom) and injected in Hank’s balanced salt solution (HBSS). Fetal livers were harvested from BubR1^H/H^ E14.5 embryos, homogenized by pipetting, passed through a 70 μM cell strainer, and frozen in FBS + 5% dimethylsulfoxide in liquid nitrogen. On the day of reconstitution, fetal liver cells were thawed in Iscove’s modified Dulbecco’s medium (IMDM) supplemented with 2% FBS. Viability was assessed by propidium iodide staining with a FACSCalibur flow cytometer (Becton Dickinson). Live cells were then counted and injected in HBSS. All BubR1^H/H^ animals were genotyped by Transnetyx, following protocols described in Baker *et al.* (2004).

Peripheral blood was collected with heparinized capillary tubes into sodium heparin diluted in PBS. Following the lysing of red blood cells in ACK lysing buffer, cells were incubated with an anti-CD45.1 antibody, obtained from Biolegend (A20), per the manufacturer’s specifications. CD45.1-negative cells were isolated via Aria I cell sorter (Beckerson Dickinson). These while blood cells were analyzed by single-cell sequencing, following the protocol in Knouse *et al.* (2014).

